# A human antibody with blocking activity to RBD proteins of multiple SARS-CoV-2 variants including B.1.351 showed potent prophylactic and therapeutic efficacy against SARS-CoV-2 in rhesus macaques

**DOI:** 10.1101/2021.02.07.429299

**Authors:** Chunyin Gu, Xiaodan Cao, Zongda Wang, Xue Hu, Yanfeng Yao, Yiwu Zhou, Peipei Liu, Xiaowu Liu, Ge Gao, Xiao Hu, Yecheng Zhang, Zhen Chen, Li Gao, Yun Peng, Fangfang Jia, Chao Shan, Li Yu, Kunpeng Liu, Nan Li, Weiwei Guo, Guoping Jiang, Juan Min, Jianjian Zhang, Lu Yang, Meng Shi, Tianquan Hou, Yanan Li, Weichen Liang, Guoqiao Lu, Congyi Yang, Yuting Wang, Kaiwen Xia, Zheng Xiao, Jianhua Xue, Xueyi Huang, Xin Chen, Haixia Ma, Donglin Song, Zhongzong Pan, Xueping Wang, Haibing Guo, Hong Liang, Zhiming Yuan, Wuxiang Guan, Su-Jun Deng

## Abstract

Severe acute respiratory syndrome coronavirus-2 (SARS-CoV-2), which causes coronavirus disease-2019 (COVID-19), interacts with the host cell receptor angiotensin-converting enzyme 2 (hACE2) via its spike 1 protein for infection. After the virus sequence was published, we identified two potent antibodies against SARS-CoV-2 RBD from antibody libraries using a phage-to-yeast (PtY) display platform in only 10 days. Our lead antibody JMB2002, now in a phase I clinical trial, showed broad-spectrum *in vitro* blocking activity against hACE2 binding to the RBD of multiple SARS-CoV-2 variants including B.1.351 that was reportedly much more resistant to neutralization by convalescent plasma, vaccine sera and some clinical stage neutralizing antibodies. Furthermore, JMB2002 has demonstrated complete prophylactic and potent therapeutic efficacy in a rhesus macaque disease model. Prophylactic and therapeutic countermeasure intervention of SARS-CoV-2 using JMB2002 would likely slow down the transmission of currently emerged SARS-CoV-2 variants and result in more efficient control of the COVID-19 pandemic.

## Introduction

Coronavirus disease-2019 (COVID-19), which is caused by the novel severe acute respiratory syndrome (SARS) coronavirus-2 (SARS-CoV-2), was first reported at the end of 2019 and has spread worldwide as a severe pandemic (1). As of February 6, 2021, the World Health Organization (https://www.who.int) has received reports of 104,956,439 confirmed cases including 2,290,488 deaths. Of much more public health concern, numerous SARS-CoV-2 mutants continue to emerge, some of which are reported to have enhanced transmissibility or reduced protective effect by vaccines (2). Such strains are South African mutant B.1.351 and UK mutant B.1.1.7, which spread faster than SARS-CoV-2 prototype and have aroused even more concerns all over the world.

To date, no typical therapies or repurposed drugs have shown the desired efficacy in treating COVID-19 (3); however, monoclonal antibodies (mAbs) are a potential effective therapeutic option. The safety and potency of human antibodies targeting viral surface proteins have been demonstrated in multiple clinical trials investigating infectious diseases, such as Ebola (4,5), SARS (6,7), and Middle East respiratory syndrome (8). Recently, the U.S. Food and Drug Administration issued an emergency use authorization for the investigational mAb therapy bamlanivimab (LY-CoV555; Lilly) and combination mAb therapy with casirivimab and imdevimab (Regeneron) for the treatment of mild-to-moderate COVID-19 (https://www.fda.gov/).

Because SARS-CoV-2 infection is initiated by attachment of the spike (S) glycoprotein on the viral surface to the angiotensin-converting enzyme 2 (ACE2) receptor on the host cell via the viral receptor-binding domain (RBD) (9), antibodies that target the RBD are hypothesized to have potent neutralizing activity. SARS-CoV-2 RBD neutralizing antibodies (nAbs) have been isolated from B cells of convalescent patients via single-cell sequencing (10–13). However, despite the ability of single-cell sequencing to quickly identify thousands of antigen-binding sequences, tremendous effort is required to produce and profile hundreds or even thousands of recombinant antibodies to obtain nAbs with the desired properties (10–13). By contrast, we applied a phage-to-yeast (PtY) platform (14) that combines the advantages of phage display (15,16) and yeast display (17) to precisely and efficiently identify the desired nAbs from naïve human B cell antibody libraries.

Accordingly, in this study, we report the rapid identification of two potent nAbs against SARS-CoV-2 from our naïve phage-displayed human B cell scFv libraries using a PtY platform. The most potent antibody, JMB2002 (Ab2001.08 N297A), not only showed potent in *vitro* blocking activity against a broad-spectrum of SARS-CoV-2 variants including the B.1.351 lineage but also potent therapeutic efficacy and complete prophylactic protection against SARS-CoV-2 in a rhesus macaque infection model.

## Results

### RBD-binding mAbs were precisely and efficiently identified with the PtY platform

The PtY platform was used to quickly identify potential clones with neutralizing activity against SARS-CoV-2 (Fig. 1). First, we screened binders with biotinylated SARS-CoV-2 RBD protein in solution using an in-house naïve phage displayed human B cell scFv library (library size: 2.8 × 10^10^) to identify mAbs. Subsequently, yeast displayed scFv libraries were constructed from enriched phage scFv display outputs with more than 10-fold coverage to maintain diversity. Specifically, we added hACE2 to a mixture of the yeast display library and the SARS-CoV-2 RBD protein to select clones with potential hACE2-neutralizing activity in solution. After sorting by FACS and sequencing, 117 potential neutralizing clones with unique sequences were identified, and the sequences of these clones were analyzed to avoid those with potential post-translational modifications, which may affect antibody function and stability. Thirty-four clones from the naïve phage display library were selected for further characterization.

**Fig. 1.**
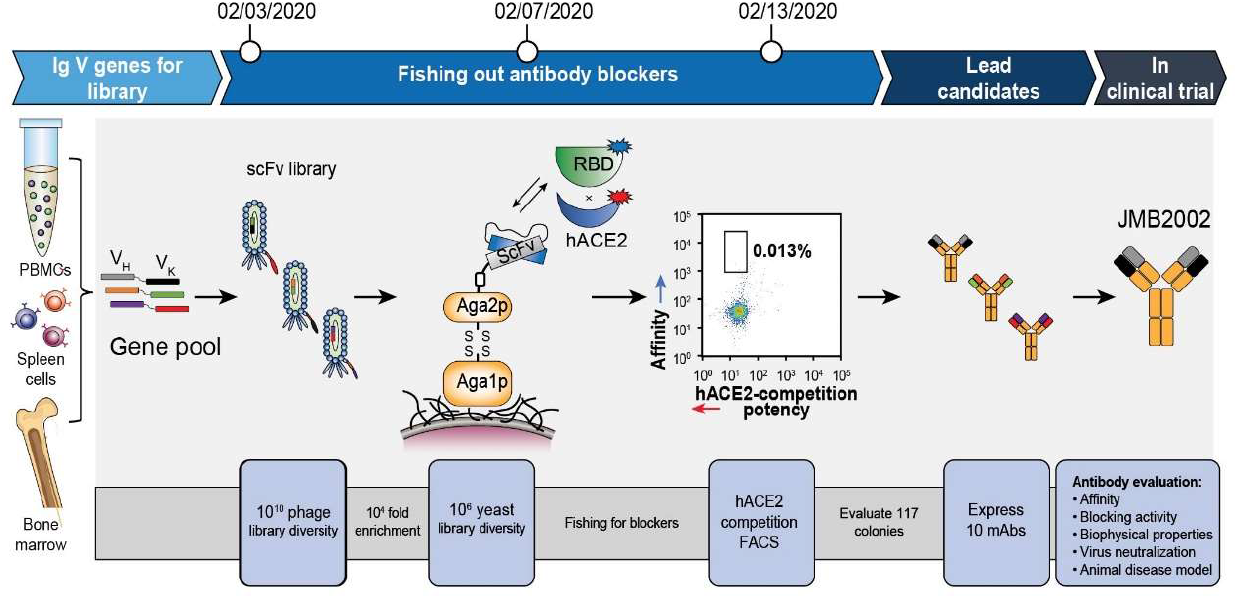
Identification of neutralizing antibodies with a PtY display platform. We first used our preconstructed naïve phage displayed human scFv library to screen binders with biotinylated SARS-CoV-2 RBD protein in the solution phase. After enrichment of phage binders, the scFv DNA from enriched binders was cloned into the yeast display plasmid, resulting in display of scFvs on the yeast cell surface. We then performed FACS to isolate potential blocking antibodies that could prevent binding of the SARS-CoV-2 RBD to hACE2. The 0.013% gate contained blocking antibodies with high affinity toward RBD. That is, higher Y axis signal represented higher affinity to labeled RBD, whereas lower X signal represented higher potency in blocking the binding of differently labeled hACE2 to RBD. The potential blocking antibodies were sent for sequencing and transient expression. The purified antibodies were evaluated for affinity, blocking activity, biophysical properties, and virus-neutralizing activity.

To confirm the specificity of these clones, their ability to bind to the SARS-CoV-2 RBD and to two nonrelated control proteins and their ability to block the binding of the CfSARS-CoV-2 RBD to hACE2 were tested. Ten clones from the naïve phage display library were identified, and the corresponding mAbs were produced in mammalian cells. We next used biolayer interferometry (BLI) to determine the affinity of selected and purified mAbs toward the SARS-CoV-2 RBD. As shown in Table 1, five of the ten antibodies from the naïve phage display library bound to the SARS-CoV-2 RBD with high affinity indicating that the PtY platform successfully identified high-affinity SARS-CoV-2 RBD-binding mAbs.

**Table 1.** Characteristics of potential blocking antibodies. NA: not applicable.

| Antibody | Expression level (mg/L) | SARS-CoV-2 RBD | SARS-CoV-2 RBD | Epitope bin | Live virus neutralization on Viral load reduction (Log<sub>10</sub>) |
| --- | --- | --- | --- | --- | --- |
|  |  | binding | blocking |  |  |
|  |  | Affinity<br>K<sub>D</sub> (nM) | IC<sub>50</sub> (nM) |  | 24 h |
| Ab2001.02 | 105 | 19.50 | 13.68 | 3 | 0.22 |
| Ab2001.03 | 116 | 10.90 | 5.85 | 3 | 0.18 |
| Ab2001.08 | 153 | 5.19 | 0.52 | 1 | 2.18 |
| Ab2001.09 | 112 | 11.90 | 3.13 | 2 | 2.10 |
| Ab2001.10 | 118 | 6.27 | 0.97 | 2 | 2.20 |
| 8 + 10 | NA | NA | 1.67 | 1 + 2 | 2.13 |
| hACE2 | NA | 37.30 | 34.80 | NA | NA |

### RBD-binding mAbs showed blocking and live virus-neutralizing activity

To evaluate the neutralization ability of the antibodies with specific and strong SARS-CoV-2 RBD-binding activity, we used enzyme-linked immunosorbent assay (ELISA) to test their ability to block SARS-CoV-2 RBD-hACE2 binding. Among the antibodies tested, Ab2001.08 and Ab2001.10 most potently blocked the binding of the SARS-CoV-2 RBD to hACE2 (Fig. 2A). Moreover, to determine whether these lead mAbs recognize the same or different RBD epitopes, an epitope binning experiment using BLI was performed, which demonstrated that Ab2001.08 and Ab2001.10 were grouped in different epitope bins (Fig. 2B). These results showed that the PtY platform precisely identified potent antibodies that blocked RBD binding to hACE2.

**Fig. 2.**
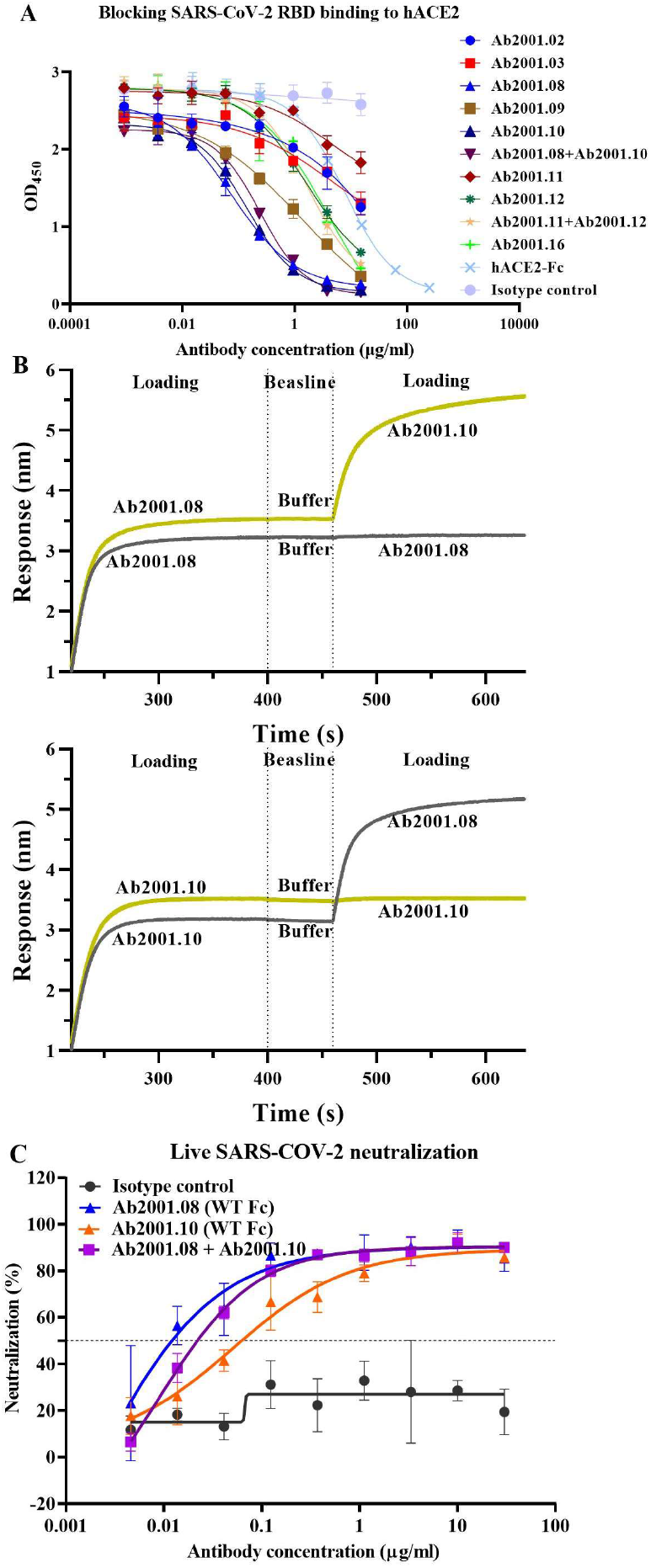
Characterization of potential blocking antibodies. **(A)** Blocking assay was performed by immobilizing 1 μg/ml hACE2 on a plate. Serially diluted antibodies and biotinylated SARS-CoV-2 RBD protein were added for competitive binding to hACE2. IC_50_ values were calculated with Prism V8.0 software using a four-parameter logistic curve fitting approach. **(B)** Epitope binning was carried out by BLI. Biotinylated SARS-CoV-2 RBD was immobilized onto the SA sensor, and a high concentration of the primary antibody was used to saturate its own binding site. Subsequently, a second antibody was applied to compete for the binding site on the SARS-CoV-2 RBD protein. Data were analyzed with Octet Data Analysis HT 11.0 software. **(C)** Neutralization activities of Ab2001.08 and Ab2001.10 were assessed by live virus assay. Live SARS-CoV-2 and serially diluted (3-fold) antibodies were added to VERO E6 cells. The PRNT_50_ values were determined by plotting the plaque number (neutralization percentage) against the log antibody concentration in Prism V8.0 software.

To measure the SARS-CoV-2-neutralizing capability of those antibodies, a live SARS-CoV-2 assay was performed by measuring the viral load 24 h after virus infection by quantitative real-time reverse transcription-PCR (qRT-PCR). Compared to an irrelevant IgG control, Ab2001.08, Ab2001.09, Ab2001.10, and Ab2001.08+Ab2001.10 resulted in a greater than 2 log reduction in the viral load at 24 h after infection (Table 1). These results indicated that these antibodies exhibited live virus-neutralizing activity.

To compare the potency of Ab2001.08 and Ab2001.10, which showed the most promising activity in inhibiting virus replication, we performed a plaque reduction neutralization tests (PRNTs) using SARS-CoV-2. Ab2001.08 and Ab2001.10 showed PRNT_50_ values of 12 and 62 ng/ml, respectively (Fig. 2C).

To select a candidate with the lowest risks for manufacture, physicochemical properties were evaluated. Ab2001.08 displayed desired CMC manufacturability (Table S1) (18) and higher virus neutralization potency than Ab2001.10. Therefore, we selected Ab2001.08 as the lead antibody for further development. Immunogenicity is also critical in the process of therapeutic antibody development (19,20). Repeated dosing of a therapeutic protein can lead to B-cell activation, which is triggered by T cell-recognition of peptide epitopes displayed on major histocompatibility complex class II (MHCII) proteins on the surface of mature antigen-presenting cells, resulting in the production of anti-drug antibodies (ADAs). ADAs can influence drug clearance and/or therapeutic efficacy. To predict potential immunogenicity risk, *in silico* analysis of antibody variable region gene sequences was performed using an in-house developed algorithm to assess potential T-cell epitopes. The algorithm provides value as a ranking tool; that is, a lower value indicates lower immunogenicity. By this algorithm, Ab2001.08 was predicted to show low immunogenicity compared to that of some marketed therapeutic antibodies (Table S2A). Additionally, JMB2002 has a comparable somatic hypermutation rate (SHM) (Table S2B) compared to nAbs isolated from convalescent COVID-19 patients, suggesting low potential immunogenicity of this antibody.

Overall, our data showed that potent nAbs with diverse epitopes and good developability could be identified by the PtY platform (Fig. 1).

### JMB2002, an antibody-dependent enhancement (ADE)-reducing mutant of Ab2001.08, exhibited good safety indicators

ADE is a side-effect of nAbs in which anti-virus antibodies enhance the entry of viruses into immune cells expressing Fc receptors (FcRs). To reduce the potential risk of ADE by reducing the interaction of the antibody with Fc receptors, we introduced the N297A mutation (21) into the Fc region of Ab2001.08, yielding Ab2001.08 N297A (JMB2002). Antibody binding to FcRs was evaluated by BLI. Ab2001.08, with wildtype human IgG1, bound to human FcγRI, FcγRIIA (R/H167), and FcγRIIIA (F/V176) with K_D_ values similar to published values (22) (Fig. 3A and Table S3). After Fc modification, JMB2002 could still bind to FcγRI (CD64), with an affinity approximately 9-fold lower than that of Ab2001.08, whereas its binding ability to FcγRIIA (R/H167) and FcγRIIIA (F/V176) was markedly reduced (Fig. 3A and Table S3).

**Fig. 3.**
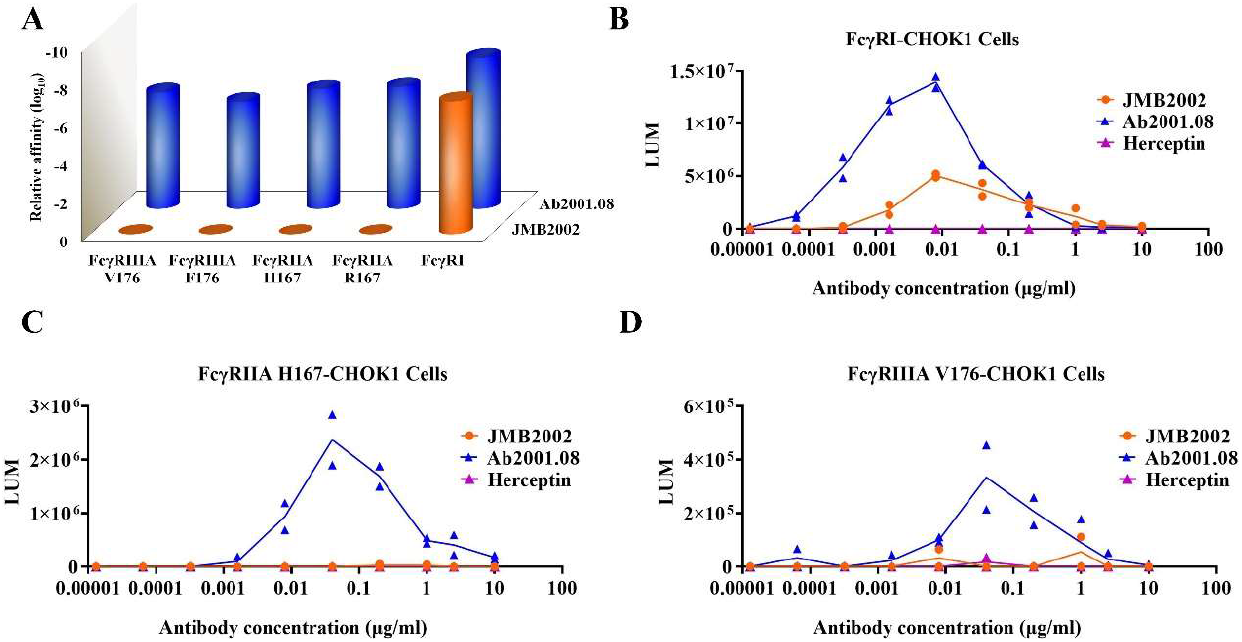
Effects of Fc modification on the ADE activity of JMB2002. **(A)** Binding of Ab2001.08 and JMB2002 to FcγRs was determined by BLI. His-tagged FcγR was loaded onto the HIS1K sensor, and serially diluted antibodies bound to the receptor on the biosensor. K_D_ values were determined with Octet Data Analysis HT 11.0 software using a 1:1 global fit model. **(B-D)** ADE activity was measured using a pseudotyped SARS-CoV-2 system containing a luciferase reporter. Pseudotyped viruses were preincubated with serially diluted antibodies for 1 h. The mixture was added to FcγR-expressing cells and incubated at 37°C for 20-28 h. Infection of cells with pseudotyped SARS-CoV-2 was assessed by measuring cell-associated luciferase activity. Herceptin was used as the irrelevant IgG control.

To evaluate ADE activity, we infected Fcγ receptor-engineered cell lines with SARS-CoV-2 pseudovirus in the presence of antibodies. Consistent with the binding affinity of the antibodies to FcγRI, Ab2001.08 mediated greater virus infection in the FcγRI-engineered cell line than did JMB2002 (Fig. 3B). As expected, no ADE effect was observed for the control antibody Herceptin. Moreover, because JMB2002 binds only weakly to FcγRIIA and FcγRIIIA, the ADE effect was abolished for JMB2002 when FcγRIIA H167- and FcγRIIIA V176-engineered cell lines were used as target cells, whereas Ab2001.08 retained its ADE activity (Fig. 3C, D). In summary, Fc-modified JMB2002 reduced ADE activity in the FcγRI-expressing cell line and eliminated the ADE effect in the FcγRIIA H167- and FcγRIIIA V176-expressing cell lines, indicating that JMB2002 may have a good safety profile *in vivo*.

### JMB2002 was a highly potent, broad-spectrum SARS-CoV-2-nAb with good manufacturability

To further study the neutralizing ability of JMB2002, we first determined its binding affinity for SARS-CoV-2 S proteins by BLI. JMB2002 bound to the SARS-CoV-2 RBD with a K_D_ of approximately 3.33 nM and to the tested SARS-CoV-2 RBD variants with a K_D_ in the single-digit nM range (Fig. 4A and Table S4A). The binding of JMB2002 to a double-mutant RBD (N354D/D364Y) and a mutant RBD (L452R) was decreased approximately 3.5- and 65-fold, respectively, compared with its binding to a prototype RBD (Fig. 4A and Table S4A). JMB2002 also bound to SARS-CoV-2 S1 (D614G), a currently widespread strain with increased infectivity, with a K_D_ of 11.9 nM. Moreover, the binding affinity was similar to that for SARS-CoV-2 S1 (Fig. 4B and Table S4B). We also assessed the binding activity of JMB2002 to recently emerged SARS-CoV-2 mutants from UK and South Africa. Encouragingly, JMB2002 showed 1.9- and 6.8- fold higher affinity to the UK and South African mutants when compared to S1 prototype, respectively (Fig. 4B and Table S4B). These results demonstrated that JMB2002 could bind to a broad range of SARS-CoV-2 RBD variants with high affinity, indicating that this mAb has broad-spectrum activity against SARS-CoV-2 strains.

**Fig. 4.**
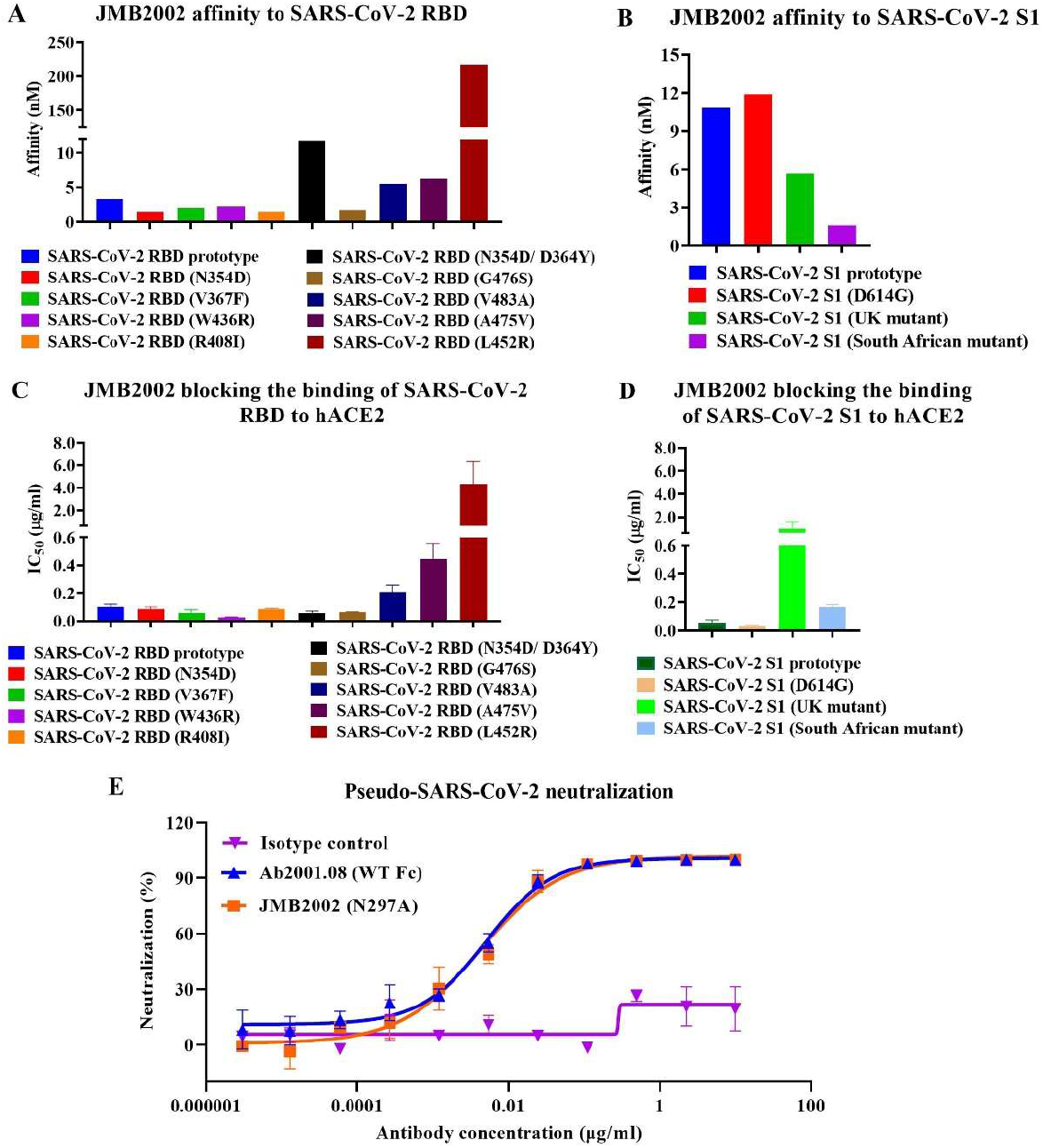
Characterization of JMB2002. Binding affinity of JMB2002 for the SARS-CoV-2 RBD (**A**)/S1 (**B**) prototype and its variants was determined by BLI. JMB2002 was loaded onto the AHC sensor, and serially diluted antigens were bound to JMB2002 on the biosensor. K_D_ values were determined with Octet Data Analysis HT 11.0 software using a 1:1 global fit model. Blocking activity was assessed using ELISA with hACE2-coated plates. A mixture of biotinylated SARS-CoV-2 RBD (**C**)/S1 (**D**) proteins and JMB2002 was added for competitive binding to hACE2. IC_50_ values were calculated by Prism V8.0 software using a four-parameter logistic curve fitting approach. Values are displayed as the mean ± standard deviations from three independent experiments. **(E)** The pseudovirus neutralization activity of JMB2002 was evaluated using a pseudotyped SARS-CoV-2 system, which contained a luciferase reporter. Pseudotyped viruses were preincubated with serially diluted antibodies for 1 h. The mixture was added to hACE2-expressing cells and incubated at 37°C for 20-28 h. Infection of cells with pseudotyped SARS-CoV-2 was assessed by measuring cell-associated luciferase activity. IC_50_ values were calculated by plotting the inhibition rate against the log antibody concentration in Prism V8.0 software.

We next performed ELISA to test the blocking activity of JMB2002. JMB2002 blocked the binding of SARS-CoV-2 S proteins to hACE2 to different extents (Fig. 4C, D and Table S4A, B). Specifically, JMB2002 displayed decreased blocking activity toward the V483A, A475V, and L452R mutant RBDs by approximately 2-, 4.4-, and 42-fold, respectively, compared to that of the RBD prototype (Fig. 4C and Table S4A). In addition, the blocking effect of JMB2002 on the N354D/D364Y and W436R RBDs was approximately 1.7- and 4.3-fold, respectively, stronger than that on the RBD prototype (Fig. 4C and Table S4A). Although hACE2-Fc prevented the SARS-CoV-2 RBD from binding to hACE2, it was approximately 67-fold less potent than JMB2002 (Table 1), indicating that JMB2002 could be much more potent than hACE2-Fc for treating COVID-19. Notably, JMB2002 demonstrated potent blocking activity against the South African mutant even at the cost of a 2.8-fold loss in potency when compared to that of S1 prototype (Fig. 4D and Table S4B). Interestingly, the blocking activity of JMB2002 toward the UK mutant was decreased 17-fold (Fig. 4D and Table S4B) although the binding activity increased by 1.9-fold (Fig. 4B and Table S4B), as compared to S1 prototype. Moreover, the potency of JMB2002 was 12-fold and 3.4- fold higher than that of hACE2-Fc against the South African and UK mutants, respectively (Fig. S1). The activity of antibody CB6, which was included as a comparator, was impaired against the UK mutant and markedly abolished against South African mutant (Fig. S1).

To determine the neutralizing activity of JMB2002, we performed a pseudovirus neutralization assay in which the entry of pseudovirus into the hACE2-expressing cell line was correlated with an increased luciferase signal in the cells. JMB2002 and Ab2001.08 showed potent neutralization ability, with half-maximal inhibitory concentration (IC_50_) values of 4.25 and 5.09 ng/ml, respectively, against SARS-CoV-2 pseudovirus (Fig. 4E). Additionally, Fc modification of Ab2001.08 had no effect on its neutralizing activity.

Based on the potent neutralizing activity and desired developability described above (Table S1), we moved JMB2002 into the CMC development stage. The expression titer of the obtained CHO-K1 CMC clone was approximately 6 g/L. Additionally, the monomer level of JMB2002 was 98.88%, as measured by size-exclusion chromatography-high performance liquid chromatography (SEC-HPLC), and the Fab Tm was 87.7°C, suggesting the good manufacturability of JMB2002. Overall, our results suggested that JMB2002 is a broad-spectrum potent nAb with desired manufacturability.

### JMB2002 showed potent prophylactic and therapeutic efficacy in rhesus macaques

To assess the ability of JMB2002 to treat viral infection and protect against viral challenge in vivo, we tested this antibody in rhesus macaques infected with SARS-CoV-2. In this experiment, rhesus macaques were assigned to three experimental conditions: the control animal was intravenously (IV) administered a single dose of irrelevant hIgG1 N297A (20 mg/kg) one day post-infection (dpi) with 1 × 10^5^ 50% tissue culture infectious dose (TCID_50_) of SARS-CoV-2; the prophylactic group was intravenously administered 20 mg/kg JMB2002 one day before viral infection; and the therapeutic group was intravenously administered 50 mg/kg JMB2002 one day and three days after viral infection (Fig. 5A). The viral load of oropharyngeal swab samples in the control group peaked at approximately 10^7^ RNA copies/ml at 5 dpi (Fig. 5B). In contrast, a single dose of JMB2002 (20 mg/kg) before infection provided complete prophylactic protection against SARS-CoV-2 infection, with no virus detected in oropharyngeal swab samples (under the detection limit of 200 copies/ml, Fig. 5B). In the therapeutic group, the viral RNA of one animal was reduced by approximately 3 logs compared with that of the control animal at 5 dpi (Fig. 5B). Notably, the viral load of the other animal in the therapeutic group was reduced to less than the detection limit at 3 dpi (Fig. 5B). These results suggested that JMB2002 can provide complete protection and potent therapeutic efficacy *in vivo*.

**Fig. 5.**
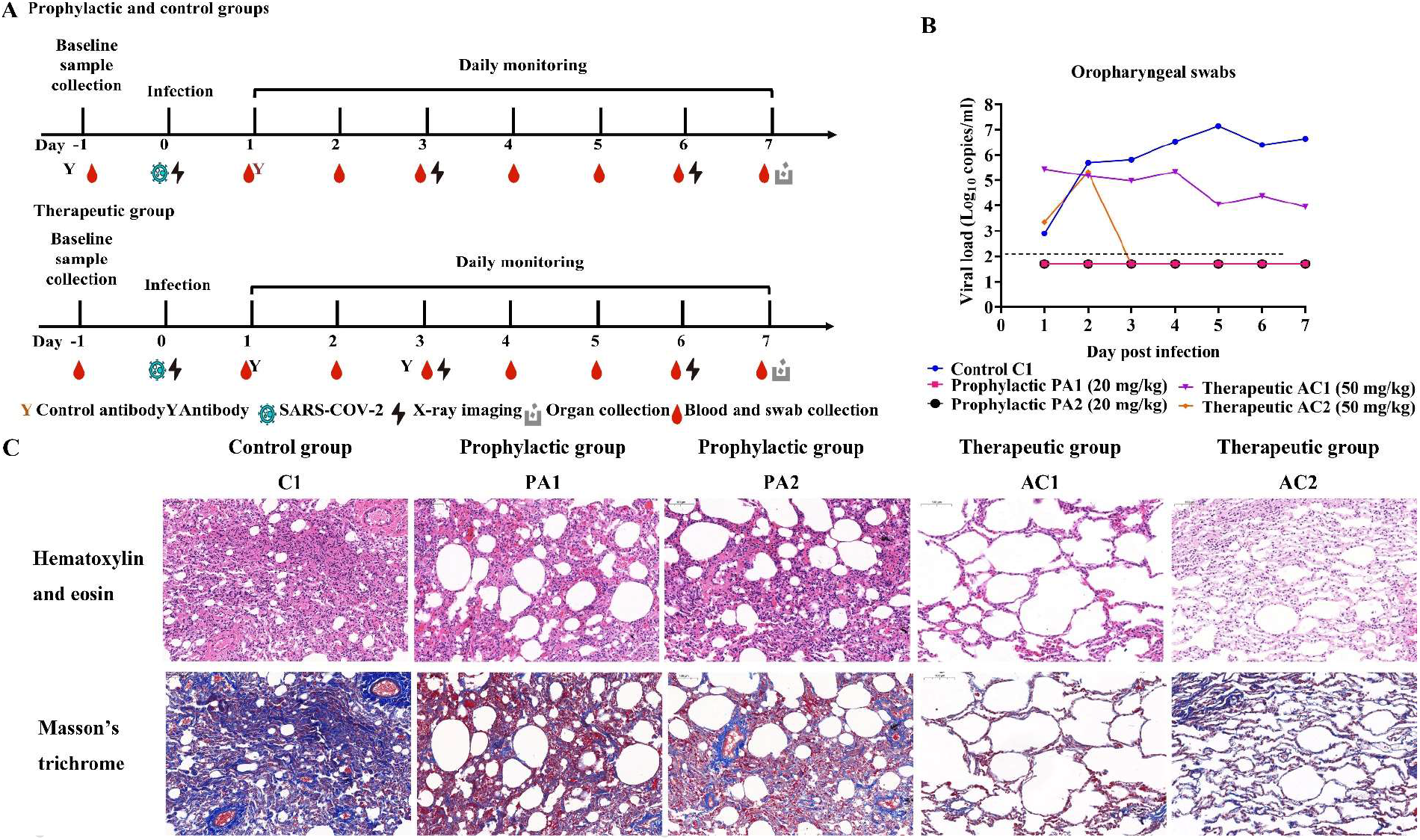
Prophylactic and therapeutic efficacies of JMB2002 against SARS-CoV-2 infection in rhesus macaques. **(A)** Schematic representation of the design of the in vivo animal experiment. Five monkeys were divided into three groups: the control group (one animal, C1), prophylactic group (two animals, PA1 and PA2), and therapeutic group (two animals, AC1 and AC2). In the prophylactic group, a single dose of 20 mg/kg JMB2002 was intravenously injected into the animals before SARS-CoV-2 infection. The next day, all monkeys were infected with virus (1 × 10^5^ TCID_50_) via intratracheal inoculation. In the therapeutic group, 50 mg/kg JMB2002 was injected at 1 and 3 dpi, whereas in the control group, a single dose of 20 mg/kg irrelevant IgG control was administered at 1 dpi. **(B)** The viral load in oropharyngeal swabs was monitored for 7 days by qRT-PCR. The dotted line indicates the copy number detection limit. **(C)** Histopathological and immunohistochemical characterization of lung tissues. All animals were euthanized and necropsied at 7 dpi. The tissue samples were collected, fixed in 10% formalin solution, embedded in paraffin, sectioned, and stained with hematoxylin and eosin or Masson’s trichrome before observation by light microscopy. Scale bar = 100 μm.

Because COVID-19 is predominantly a lung infection, we also examined the ability of JMB2002 to protect against lung damage from SARS-CoV-2 infection. All animals were sacrificed and necropsied at 7 dpi. Based on evaluation of pneumonia severity, the control animal was diagnosed with viral pneumonia with extensive pulmonary fibrosis, exhibiting pathology including thickened alveolar septa, fibroblast proliferation and fibrosis, and intensive monocyte and lymphocyte infiltration (Fig. 5C). In some alveolar cavities, cellulose exudation was observed, with hyaline membrane formation and pulmonary hemorrhage. By contrast, animals treated with JMB2002 before or after infection displayed very limited pathological lung damage characterized by an overall intact alveolar structure, reduced edema and the absence of hyaline membrane formation, with less fibrosis and less leukocyte infiltration than the control animal (Fig. 5C). In summary, JMB2002 reduced infection-related lung damage in both the prophylactic and therapeutic animal groups, suggesting that JMB2002 is an effective therapeutic.

## Discussion

To rapidly develop therapeutic antibodies in response to SARS-CoV-2 infection, we applied a PtY platform to identify a potent nAb JMB2002 in a precise and efficient manner, suggesting that the PtY platform can meet the needs of an immediate response to a global pandemic by accelerating research and development of nAbs. PtY, a unique human mAb discovery platform, offers the following 5 advantages: 1) the ability to identify lead SARS-CoV-2 nAbs from a naïve scFv library without using B cells from COVID-19 convalescent patients; 2) the ability to isolate rare nAbs from the libraries containing over 10^10^ antibody clones within 10 days; 3) the ability to maintain the native conformation of the RBD, antibodies, and hACE2 at their maximal levels, because all screens were performed in solution with the inclusion of hACE2 as a competitor, leading to maximum possible simulation of hACE2 by the antibody; 4) the ability to identify two highly potent mAbs with different epitopes by producing and profiling only 10 IgGs; and 5) the ability to monitor antibody expression levels in real time, an important criterion in CMC. In fact, the expression titer of JMB2002 was approximately 6 g/L in the CHO-K1 CMC clone. Based on published literature (10–13), SARS-CoV-2 nAbs identified from B cells of convalescent COVID-19 patients via single-cell technologies often require the production and profiling of hundreds to thousands of antibodies, highlighting the advantages of the PtY approach.

ADE has been observed in dengue virus infection (23,24), and several reports have demonstrated that antibodies induced by the SARS-CoV S protein enhanced viral entry into Fc receptor-expressing cells (25,26). To mitigate the potential risk of ADE, the N297A mutation was introduced into the Fc of JMB2002. Neutralization assays with pseudotyped SARS-CoV-2 confirmed the high potency of JMB2002, and Fc modification had no effect on its in *vitro* neutralizing activity. JMB2002 also displayed a complete protective effect and potent therapeutic efficacy in an in vivo nonhuman primate model, with substantially reduced lung tissue damage, suggesting that JMB2002 may have maintained a balance between attenuated Fc-mediated effector function such as antibody-dependent cellular phagocytosis (ADCP) based viral clearance efficacy and reducing potential ADE risk. Different Fc modification strategies that involve either attenuating (13) or enhancing Fc effector function (27,28) have also been applied in the design of SARS-CoV-2 nAbs, some of which have moved into clinal trials. Our prospective goal is to have JMB2002 approved for the treatment of COVID-19, which will contribute not only to the treatment of SARS-CoV-2 infection but also to the expansion of knowledge on the impact of Fc modification on the efficacy of antibody treatment of viral infection.

The lead antibody showed broad-spectrum activity against a range of SARS-CoV-2 RBD variants. It is generally accepted that mutations in amino acids of viral surface proteins can change the viral infectivity and/or reactivity of nAbs. A total of 329 naturally occurring SARS-CoV-2 S protein mutations have been reported worldwide (29,30), which has aroused interest concerning the impact of these variants on the COVID-19 pandemic and treatment approaches. A single D614G mutation outside the RBD region of the SARS-CoV-2 S protein, the predominant mutation worldwide, has been shown to confer increased infectivity in several studies (31–33). Notably, our identified antibody JMB2002 not only bound to the SARS-CoV-2 S1 prototype and SARS-CoV-2 S1 (D614G) mutant with similar affinity but also exhibited improved potency in blocking the binding of SARS-CoV-2 S1 (D614G) to hACE2. Furthermore, three mutations, i.e., L452R, A475V, and V483A, have been shown to exhibit marked resistance to some antibodies (29). Specifically, the neutralizing activity of antibodies B38, P2C-1F11, CB6, and 247 against A475V was decreased approximately 100-fold. Moreover, RBD L452R also reduced the sensitivity to antibodies X593 and P2B-2F6 and V483A to antibody X593 by approximately 100-fold (29). In contrast, JMB2002 only slightly decreased blocking ability for V483A and A475V compared to the RBD prototype. Similar to antibodies X593 and P2B-2F6 (29), JMB2002 also displayed resistance to L452R. Furthermore, JMB2002 exhibited increased binding to N354D/D364Y and W436R. The recent emergence of new variants from UK (B.1.1.7 lineage) (34) and South Africa (B. 1.351 lineage) (35) has attracted increasing attention, because impaired efficacy of current nAb therapies or vaccines caused by these new variants was reported (36). The tested nAbs which are in clinical use or under clinical investigation showed comparable or slightly impaired efficacy against UK variant when compared to D614G mutant (36). Notably, the neutralizing activity of REGN10933, LY-CoV555 or the combination of LY-CoV555 and CB6 against South African variant was markedly or completely abolished. JMB2002 retained potent blocking activity toward South African mutant, whereas the potency against the UK mutant was 17-fold lower when compared to S1 prototype. Moreover, the potency of JMB2002 was 12-fold and 3.4-fold higher than that of hACE2-Fc against the South African and UK mutants, respectively, further indicating effective blocking of these two mutants by JMB2002. Collectively, these findings suggested the possibility of hACE2 receptor mimicry by JMB2002, indicating the potential ability of JMB2002 to exhibit broad-spectrum blocking activity against RBD mutants, an intuitive conclusion, since JMB2002 was identified by its ability to compete for hACE2 binding in solution.

Encouraged by the successful clinical trial results of the anti-SARS-CoV-2 antibody cocktail from Regeneron and inspired by their studies suggesting that treatment with a combination of antibodies targeting non-competing epitopes prevented virus escape (37), attempts to optimize the sequence of Ab2001.10 to increase its binding affinity for potential therapeutic application in combination with JMB2002 could be considered, since it recognizes different epitope from JMB2002. Bispecific antibodies constitute an even more attractive approach than combination/cocktail therapies because of shorter development time and lower manufacturing cost.

In this study, we report the rapid discovery and development of JMB2002 into clinical trial stage. JMB2002 has demonstrated complete prophylactic and potent therapeutic efficacy in a rhesus macaque disease model. JMB2002 showed broadspectrum in *vitro* neutralization activity toward multiple SARS-CoV-2 mutants including D614G, UK and South African variants. Therefore, this antibody, alone or in combination with other therapeutics, offers a potential therapy to prevent virus escape and end current COVID-19 pandemic earlier.

## Methods

### Monospecific antibodies identified by the PtY platform

To obtain SARS-CoV-2 RBD-binding clones, we screened preconstructed human naïve phage scFv libraries with biotinylated SARS-CoV-2 RBD (Sino Biological, 40592-V05H). Phage libraries were constructed using antibody gene fragments amplified from PBMCs of 50 healthy human subjects (Allcells, PB003F and PB003C) and total RNA from PBMCs (TaKaRa, 636592), spleens (TaKaRa, 636525), and bone marrow (TaKaRa, 636591) from 494 healthy human subjects. After two rounds of phage selection to enrich for SARS-CoV-2 RBD binders, the scFv DNA from phagemids was amplified and cotransferred with the yeast display plasmid into the *Saccharomyces cerevisiae* strain EBY100 (ATCC, MYA-4941) via electroporation, resulting in a yeast display scFv library (38).

We then developed a novel competitive FACS approach to rapidly identify potential blocking antibodies (Fig. 1). Specifically, the library were incubated with SARS-CoV-2 RBD containing a mouse Fc tag (Sino Biological, 40592-V05H), and biotinylated hACE2 (Kactus, ACE-HM401) was then added. Next, the libraries were labeled with streptavidin (SA)-phycoerythrin (PE) (eBioscience, 12-4317-8) and goat anti-mouse-Alexa Fluor 647 antibodies (Thermo Fisher Scientific, A-21235). The population with a weak PE signal and a strong Alexa Fluor 647 signal was sorted in a FACSAria II (BD) to identify potential blocking antibodies with high RBD binding activity. The collected cells were sent for sequencing and further colony evaluation. The colonies with unique sequences were labeled with SA-PE and goat anti-mouse-Alexa Fluor 647 antibodies to confirm the hACE2 blocking potency. The colonies were also labeled with a biotinylated mouse Fc containing irrelevant antigen to confirm specific binding.

### Transient expression of monospecific IgG antibodies

The full-length sequences of human IgG1 light and heavy chains of the candidate antibodies were codon optimized, synthesized, and cloned into an expression vector (GenScript). The antibody production was conducted by co-transfection of the light chain and heavy chain sequences into Expi293 cells according to the manufacturer’s instructions (Thermo Fisher Scientific, A14635). The culture supernatant was collected at 7 days post-transfection, and protein purification was performed with Protein A magnetic beads (GenScript, L00695). The purified mAbs were dialyzed against phosphate-buffered saline (PBS) and kept for further analysis of biophysical properties and biological activities.

### Measurement of antibody affinity by BLI

The affinity of mAbs for SARS-CoV-2 RBD/S1 and its mutants (SARS-CoV-2 RBD [ACRO, SPD-C52H3], SARS-CoV-2 S1 [ACRO, S1N-C52H4], SARS-CoV-2 RBD [N354D/D364Y] [ACRO, SPD-S52H3], SARS-CoV-2 RBD [V367F] [ACRO, SPD-S52H4], SARS-CoV-2 RBD [N354D] [ACRO, SPD-S52H5], SARS-CoV-2 RBD [W436R] [ACRO, SPD-S52H7], SARS-CoV-2 RBD [R408I] [ACRO, SPD-S52H8], SARS-CoV-2 RBD [G476S] [ACRO, SPD-C52H4], SARS-CoV-2 RBD [V483A] [ACRO, SPD-C52H5], SARS-CoV-2 RBD [A475V] [ACRO, SPD-C52Hd], SARS-CoV-2 RBD [L452R] [ACRO, SPD-C52He], SARS-CoV-2 S1 [D614G] [ACRO, S1N-C5256], SARS-CoV-2 S1 [K417N, E484K, N501Y, D614G] [Sino Biological, 40591-V08H10], SARS-CoV-2 S1 [HV69-70 deletion, Y144 deletion, N501Y, A570D, D614G, P681H] [Sino Biological, 40591-V08H12]) was measured using Octet Red96 (ForteBio, Sartorius). We used anti-human Fc (AHC) biosensors (ForteBio, 18-5060) to load different mAbs (5 μg/ml) in kinetics buffer (0.02% Tween-20, 0.1% bovine serum albumin [BSA] in PBS) for 40 s. After immersion for 3 min in kinetics buffer, we incubated the mAb-coated sensors with different concentrations of SARS-CoV-2 proteins and recorded association curves for 3 min. Then, we transferred the sensors to wells containing kinetics buffer and recorded dissociation for 10 min. We calculated K_D_ values with Octet Data Analysis HT 11.0 software using a 1:1 global fit model.

Additionally, we determined the affinity of mAbs for human Fc gamma receptors (FcγRs, including FcγRI [ACRO, FCA-H5251], FcγRIIa R167 [ACRO, CDA-H5221]/H167 [ACRO, CD1-H5223], and FcγRIIIa F176 [ACRO, CDA-H5220]/V176 [ACRO, CD8-H5224]) by using anti-Penta-HIS (HIS1K) biosensors (ForteBio, 18-5120) to load His-tagged human FcγRs (5 μg/ml) in kinetics buffer for 20 s. Association of mAbs with FcγRs (750, 375, 187.5, 93.75, 46.88, 23.44, 11.72 μg/ml) was evaluated in kinetics buffer for 3 min, except for association with FcγRI, which was evaluated in kinetics buffer at 37.5, 18.75, 9.38, 4.69, 2.34, 1.17, and 0.59 μg/ml for 3 min. Dissociation in kinetics buffer was measured for 10 min. K_D_ values were calculated with Octet Data Analysis HT 11.0 software using a 1:1 global fit model.

### Evaluation of blockade of SARS-CoV-2 RBD binding to hACE2 by ELISA

The blocking assay was performed using ELISA. Specifically, 1 μg/ml purified hACE2 protein (Kactus, ACE-HM501) was coated onto a 96-well ELISA plate (Thermo Fisher Scientific, 44-2404-21) at 4°C overnight. The next day, the plates were washed with washing buffer (PBS containing 0.05% Tween-20) and blocked with blocking buffer (2% BSA [Bovogen, BSAS 1.0] in washing buffer) at 37°C for 1 h. Four-fold serial dilutions of antibodies were preincubated with an equal volume of 0.5 μg/ml biotinylated SARS-CoV-2 RBD protein for 1 h at 37°C. The antigen/antibody mixtures were added to the plates and incubated at 37°C for 1 h. Subsequently, the plates were washed and incubated with 5000-fold diluted SA-horseradish peroxidase (Sigma, S2438). Tetramethylbenzidine substrate (Biopanda, TMB-S-003) was added for color development. The absorbance was measured at 450 nm using a spectrophotometer (Spectramax M3, Molecular Devices). IC_50_ values were determined with Prism V8.0 software (GraphPad) using a four-parameter logistic curve fitting approach.

### Epitope binning by BLI

We conducted epitope binning of mAbs with an Octet Red96 (ForteBio, Sartorius) using an in-tandem format, in which mAbs competed against one another were in a pairwise combinatorial manner for binding to the SARS-CoV-2 RBD protein. Assays were performed at 30°C with continuous agitation at 1000 rpm. After obtaining an initial baseline in kinetics buffer, 1 μg/ml biotinylated SARS-CoV-2 RBD (Kactus, COV-VM4BD) was captured onto a SA biosensor (ForteBio, 18-5020) for 65 s. To saturate the binding sites for the first mAb on the SARS-CoV-2 RBD antigen, we exposed all sensors for 3 min to the first mAb (15 μg/ml) after immersion in kinetics buffer for 30 s. Then, we immersed the biosensors in kinetics buffer for 30 s and further immersed in wells containing a solution of the second mAb (15 μg/ml) for 3 min. Data analysis was performed with Octet Data Analysis HT 11.0 software using Epitope Binning mode.

### SARS-CoV-2 neutralization assay

The SARS-CoV-2 strain WIV04 (GenBank: MN996528.1) was obtained from the Wuhan Institute of Virology, Chinese Academy of Sciences. Vero E6 cells (ATCC, CRL-1586) were seeded in a 24-well plate (10^5^ cells/well) and incubated at 37°C, in 5% CO_2_ for 16 h. Subsequently, we added different mAbs (10 μg/ml) and then infected cells with SARS-CoV-2 at a multiplicity of infection of 0.005. Finally, we collected the cell culture supernatants after 24 h of infection for viral RNA extraction with a QIAamp 96 Virus QIAcube HT Kit (Qiagen, 57731) and for viral RNA copy number detection in a CFX96 Touch Real-Time PCR Detection System (Bio-Rad Laboratories). Data were processed using GraphPad Prism software (V8.0).

After the above initial screening for the ranking of 10 samples, including two CLC mAbs, one bsAb, and two antibody combinations, based on reduced viral RNA copy numbers, PRNT assay was further performed to characterize the neutralizing activities of the most potent antibodies. We mixed 3-fold serially diluted mAbs (100 μl/well, starting from 30 μg/ml) and SARS-CoV-2 (WIV04) (100 μl/well, 2000 PFU/ml) into 96-well plates and incubated the mixtures for 30 min at 37°C. A well containing only the antibody was set up as an untreated control. Then, the antibody/virus mixture was added to a monolayer of Vero E6 cells and incubated at 37°C for 1 h. The plaques were counted after 96 h of viral infection. We determined the PRNT_50_ values using a four-parameter logistic curve fitting approach (GraphPad Prism software V8.0).

### Biophysical characterization of monospecific IgG antibodies

SEC-HPLC was performed to quantify the monomer level of antibodies in a Waters Alliance 2695 HPLC system with a TOSOH TSKgel G3000W_XL_ column (300 × 7.8 mm, 5 μm). The mobile phase was 200 mM sodium phosphate (pH 6.8), and the flow rate was 0.5 ml/min. Samples were assessed by measuring the UV absorbance at a wavelength of 280 nm. Data were analyzed with Waters Empower 3 Enterprise software (Waters, MA, USA).

Hydrophobic interaction chromatography (HIC)-HPLC was performed to evaluate the hydrophobicity of antibodies in a Waters Alliance 2695 HPLC system with a Thermo MAbPacHIC-10 column (4 × 250 mm, 5 μm). Mobile phase A consisted of 1 M ammonium sulfate with 50 mM sodium phosphate (pH 7.0), and mobile phase B consisted of 50 mM sodium phosphate (pH 7.0). Samples were injected and eluted with a linear gradient at a flow rate of 0.8 ml/min and the UV absorbance was measured at a wavelength of 214 nm. Data were analyzed with Waters Empower 3 Enterprise software. A shorter retention time indicated that the antibody had increased hydrophilicity. We used the retention times of Omalizumab (14.2 min) and Tecentriq (26.8 min) as our standards, and the acceptance criterion was a retention time of less than 27 min.

The pI and charge variants were characterized by imaged capillary isoelectric focusing (iCIEF) in an iCE3 system (ProteinSimple, CA, USA). Samples were mixed with pharmalytes, 1% methyl cellulose, a pI marker and ddH_2_O to a final protein concentration of 0.2 mg/ml. Samples were pre-focused at 1500 V for 1 min and focused at 3000 V for 10 min, with detection at 280 nm. Data were acquired and analyzed with Chrom Perfect software (ProteinSimple).

A Protein Thermal Shift Dye Kit (Thermo Fisher Scientific) was applied to evaluate the Tm values of the candidates. According to the manufacturer’s instructions, samples were mixed with Protein Thermal Shift buffer and Protein Thermal Shift dye solution. Protein melt curves were run in an Applied Biosystems Real-Time PCR System (QuantStudio 3, Thermo Fisher Scientific, USA). Data were collected and imported into Protein Thermal Shift Software (Version 1.3) to calculate the Tm value of the candidates from the melt curves.

### ADE assay

The ADE effect of mAbs was measured by *in vitro* enhancement of infection by pseudotyped SARS-CoV-2 (National Institutes for Food and Drug Control, China) in CHO-K1 cells overexpressing FcγRI, FcγRIIA (H167) and FcγRIIIA (V176) (GenScript, M00588, M00598, and M00597). In brief, engineered CHO-K1 cells (100 μl/well) were seeded into 96-well plates at 5 × 10^5^ cells/ml. Subsequently, pseudovirus (650 TCID_50_/well) and serial dilutions of mAbs (from 10 to 0.000013 μg/ml at fourfold dilutions) were preincubated for 1 h and then used to infect cultured cells at 37°C for 28 h. Herceptin (Roche) was used as the irrelevant IgG control. Infection of cells was evaluated by luciferase expression, as determined with a PE Britelite Plus Assay (PerkinElmer, 6066769), and the results were read on a luminometer (PerkinElmer). Enhancement of viral infection was evaluated by plotting the luciferase activity versus the antibody concentration in GraphPad Prism software (V8.0).

### Pseudovirus neutralization assay

In the pseudovirus neutralization assay, serial dilutions of antibodies (from 10 to 0.000003 μg/ml at 4.5-fold dilutions) were preincubated with an equal volume of pseudovirus (650 TCID_50_/well) for 1 h at room temperature. Subsequently, CHO-K1 cells stably expressing hACE2 (WuXi Biologics) were seeded in 96-well plates (5 × 10^4^ cells/well), treated with the pseudovirus and antibody mixtures, and incubated at 37°C for 20-28 h. Luciferase activity was measured using PE Britelite-Plus assay reagent (PerkinElmer, 6066769). The neutralization inhibition rate was calculated with the following formula.

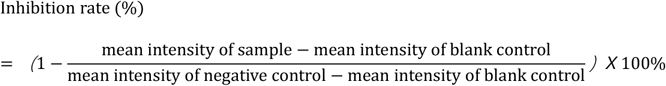

IC_50_ values were calculated by a four-parameter logistic curve fitting approach in Prism V8.0 software (GraphPad).

### *In vivo* animal model study

Five 6- to 7-year-old rhesus macaques purchased from Topgene Biotechnology (Hubei, China) were housed and cared for in laboratory animal care-accredited facility. All animal experiments were approved by the Institutional Animal Care and Use Committee of Wuhan Institute of Virology, Chinese Academy of Sciences (Ethics no. WIVA42202006). All animals were anesthetized prior to sample collection, and experiments were carried out in the animal biosafety level 4 (ABSL4) laboratory.

All animals were randomly divided into three groups: the control group (one animal, C1), prophylactic group (two animals, PA1 and PA2), and therapeutic group (two animals, AC1 and AC2). PA1 and PA2 were injected intravenously with JMB2002 (20 mg/kg) one day before infection. Subsequently, all animals were inoculated intratracheally with SARS-CoV-2 (WIV04) at a dose of 1 × 10^5^ TCID_50_. C1 was injected intravenously with irrelevant IgG (20 mg/kg) one day after infection, whereas AC1 and AC2 were injected with JMB2002 (50 mg/kg) one day and three days after infection. All animals were monitored along the timeline for recording of clinical signs and collection of oropharyngeal swabs. The animal experiment and sampling schedule are detailed in Fig. 5a.

SARS-CoV-2 RNA was detected in swab samples by qRT-PCR. Total RNA was extracted with a QIAamp Viral RNA Mini Kit (Qiagen, 52906), and qRT-PCR was conducted with a HiScript^®^ II One Step qRT-PCR SYBR^®^ Green Kit (Vazyme, Q221-01). The qRT-PCR was performed in an ABI Real-time PCR System (Applied Biosystems). Amplification was carried out with the following thermal cycling program: 50°C for 3 min, 95°C for 30 s, and 40 cycles at 95°C for 10 s and 60°C for 30 s. The qRT-PCR primer pairs targeting the SARS-CoV-2 RBD were as follows: forward primer, 5’-CAATGGTTAAGGCAGG-3’; reverse primer, 5’-CTCAAGGTCTGGATCACG-3’.

### Histopathology

Animal necropsies were performed according to a standard protocol at ABSL4. The animals were euthanized on day 7 post infection. The respiratory samples from each animal were collected for the histological analysis. Samples for histological examination were stored in 10% neutral-buffered formalin for 7 days, embedded in paraffin, sectioned and stained with hematoxylin and eosin or Masson’s trichrome prior to examination by light microscopy. A researcher with more than 10 years’ experience in the field analyzed the slides.

## Acknowledgments

We thank all colleagues from the National Biosafety Laboratory (Wuhan), Chinese Academy of Sciences, for their support during the study. We also thank our colleagues from the Center for Instrumental Rectalysis and Metrology, Wuhan Institute of Virology, Chinese Academy of Sciences. We thank Dr. C. Roger MacKenzie for critical reading and editing of our manuscript.

## Author contributions

S. D. conceived the project. C. G. and Z. W. identified the lead antibodies with the help of L. G., K. X., and Y. W. M. S. and T. H. performed the antibody production. X. L. developed an algorithm to assess potential T-cell epitopes. X. C. designed the experiment and F. J., L. Y., N. L., G. J., J. Z., and L. Y. conducted the activity assay. P. L. designed the experiment and Y. L., W. L., G. L., and C. Y. performed the biophysical analysis of antibodies. Z. X., J. X., X. H., and X. C. contributed to antibody expression experiments. Y. Y., C. S., H. M., D. S., Z. Y., and W. G. designed the live virus neutralization and in vivo animal experiments and analyzed the data. X. H., Y. Y., G. G., X. H., Y. Z., Z. C., Y. P., K. L., W. G., and J. M. performed the live virus neutralization and in vivo animal experiments. Y. Z. analyzed the HE slides. X. C. drafted the original manuscript. Z. P., X. W., H. G., H. L., Z. Y., W. G., and S. D. reviewed and edited the manuscript.

## Competing interests

C. G., X. C., Z. W., P. L., X. L., L. G., F. J., L. Y., N. L., G. J., J. Z., L. Y, M. S., T. H., Y L., W. L., G. L., C. Y, Y W., K. X., Z. X., J. X., X. H., X. C., Z. P., X. W., H. G., H. L., and S. D. were employed by Shanghai Jemincare Pharmaceuticals Co., Ltd. All other authors declared no competing interests.

## Data availability

The main data supporting the results in this study are available within the paper and Supplementary Information. Datasets are available from the corresponding authors upon reasonable request.

## Notes

### Summary of Updates

Author list updated.

