## Supplementary material for "A human antibody with blocking activity to RBD proteins of multiple SARS-CoV-2 variants including B.1.351 showed potent prophylactic and therapeutic efficacy against SARS-CoV-2 in rhesus macaques": Fig. S1

JM 036127

**This PDF file includes:**

Fig. S1
Tables S1 to S4
References (39)

**Fig. S1.**

A Blocking SARS-CoV-2 S1 (HV69-70 deletion, Y144 deletion, N501Y,A570D, D614G, P681H) binding to hACE2

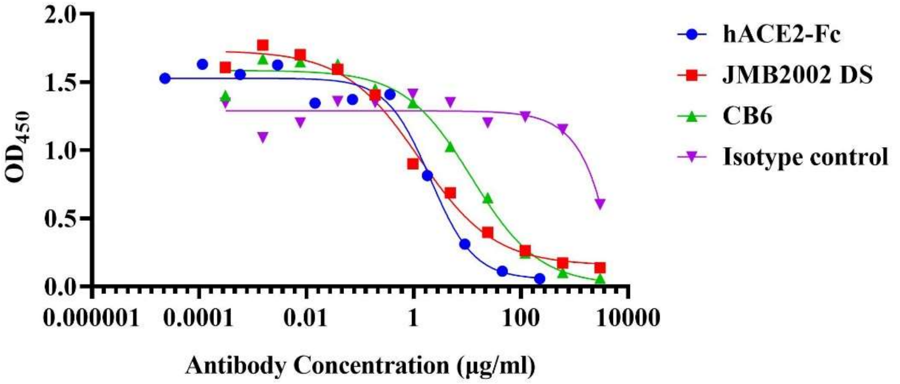


| hACE2-Fc | | JMB2002 DS | | CB6 | |
|---|---|---|---|---|---|
| mean concentration(μg/ml) | SD | mean concentration(μg/ml) | SD | mean concentration(μg/ml) | SD |
| 3.368 | 2.329 | 0.995 | 0.615 | 12.335 | 4.903 |

B Blocking SARS-CoV-2 S1 (K417N, E484K, N501Y, D614G) binding to hACE2

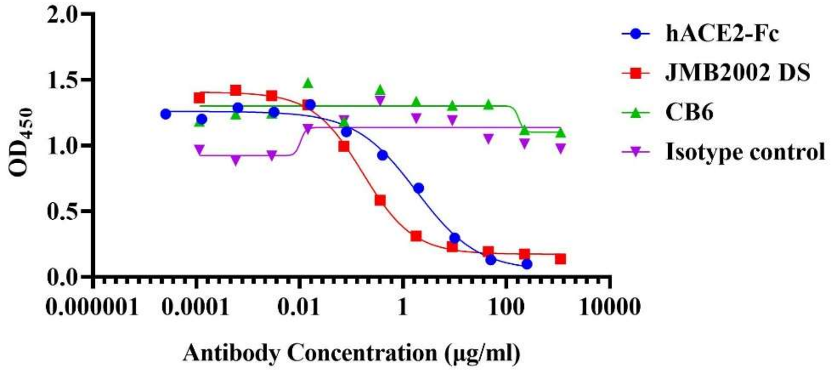


| hACE2-Fc | | JMB2002 DS | | CB6 | |
|---|---|---|---|---|---|
| mean concentration(μg/ml) | SD | mean concentration(μg/ml) | SD | mean concentration(μg/ml) | SD |
| 1.94 | 0.195 | 0.165 | 0.018 | - | - |

**Fig. S1.** Blocking activity of JMB2002 to the UK (SARS-CoV-2 S1 [H69del, V70del, Y144del, N501Y, A570D, D614G, P681H]) (A) and South African mutants (SARS-CoV-2 S1 [K417N, E484K, N501Y, D614G]) (B) was evaluated using ELISA with hACE2-coated plates. A mixture of biotinylated SARS-CoV-2 S1 proteins (1 μg/ml) and 5-fold serially diluted antibodies or hACE2 was added for competitive binding to hACE2. The $IC_{50}$ was calculated with Prism V8.0 software using a four-parameter logistic curve fitting approach. One representative figure from three independent experiments is shown.

JM 036127 JM 036127 JM 036127
