## Supplementary material for "A human antibody with blocking activity to RBD proteins of multiple SARS-CoV-2 variants including B.1.351 showed potent prophylactic and therapeutic efficacy against SARS-CoV-2 in rhesus macaques": Table S1

JM 036127

**This PDF file includes:**

Fig. S1
Tables S1 to S4
References (39)

**Table S1.**

| Antibody | Purity by SEC-HPLC (monomer %) | Fab Tm (℃) | Hydrophobicity (min) | pI value | Charge variants by iCIEF | | |
|---|---|---|---|---|---|---|---|
| | | | | | Acidic peaks (%) | Main peak (%) | Basic peaks (%) |
| Ab2001.08 | 99.5 | 87.9 | 21.6 | 7.5 | 20.8 | 76.8 | 2.4 |
| Ab2001.10 | 98.6 | 81.9 | 14.1 | 8.7 | 15.2 | 80.5 | 4.4 |

**Table S1.** Biophysical properties of Ab2001.08 and Ab2001.10

JM 036127
