## Supplementary material for "A human antibody with blocking activity to RBD proteins of multiple SARS-CoV-2 variants including B.1.351 showed potent prophylactic and therapeutic efficacy against SARS-CoV-2 in rhesus macaques": Table S2

**This PDF file includes:**

Fig. S1
Tables S1 to S4
References (39)

**Table S2A.**

| Antibody | Immunogenicity score |
|---|---|
| **Pembrolizumab** | 35 |
| **Ab2001.10** | 63 |
| **Omalizumab** | 76 |
| **Sintilimab** | 79 |
| **Ab2001.08 (JMB2002)** | 85 |
| **Trastuzumab** | 92 |
| **Nivolumab** | 105 |
| **Imdevimab (REGN10987)** | 113 |
| **Dacetuzumab** | 150 |
| **Adalimumab** | 160 |
| **Roledumab** | 173 |
| **Lorvotuzumab** | 213 |

**Table S2A.** Prediction of the immunogenicity of antibodies. *In silico* analysis of antibody variable region gene sequences was performed using an in-house algorithm to assess potential T cell epitopes. The variable region gene sequences of marketed antibodies were retrieved from the IMGT database. The variable region gene sequences of reference antibodies CB6 (PBD accession number: 7C01) and Imdevimab (PBD accession number: 6XDG) were retrieved from the PDB database.

JM 036127

**Table S2B.**

| Antibody | V-H allele | J-H allele | CDR3 length (aa) | SHM (%) |
|---|---|---|---|---|
| JMB2002 $V_\kappa$ | IGKV1-33*01 | IGKJ4*01 | 9 | 4.7 |
| JMB2002 $V_H$ | IGHV1-69*18 | IGHJ5*02 | 16 | 0.0 |
| CB6 $V_\kappa$ | IGKV1-39*01 | IGKJ2*01 | 11 | 1.8 |
| CB6 $V_H$ | IGHV3-66*01 | IGHJ4*01 | 13 | 3.4 |
| Imdevimab (REGN10987) $V_\lambda$ | IGLV2-14*03 | IGLJ3*02 | 10 | 4.5 |
| Imdevimab (REGN10987) $V_H$ | IGHV3-30*01 | IGHJ4*01 | 13 | 2.5 |

**Table S2B.** Germline analysis of antibodies.
IMGT/V-QUEST and Domain Gap Align (39) were applied to analyze antibody gene germline, complementarity determining region (CDR)3 length, and somatic hypermutation (SHM). The CDR3 length was calculated from amino acid sequences (IMGT unique numbering). The SHM frequency was calculated from the mutated amino acids. The variable region gene sequences of reference antibodies CB6 (PBD accession number: 7C01) and Imdevimab (PBD accession number: 6XDG) were retrieved from the PDB database.

JM 036127
