## Supplementary material for "A human antibody with blocking activity to RBD proteins of multiple SARS-CoV-2 variants including B.1.351 showed potent prophylactic and therapeutic efficacy against SARS-CoV-2 in rhesus macaques": Table S3

**This PDF file includes:**

Fig. S1
Tables S1 to S4
References (39)

JM 036127

**Table S3.**

| Antibody | Human FcγRs | Affinity $K_D$ (nM) |
|---|---|---|
| **Ab2001.08** | FcγRI | 11.1 |
| | FcγRIIA R167 | 373 |
| | FcγRIIA H167 | 489 |
| | FcγRIIIA F176 | 2330 |
| | FcγRIIIA V176 | 742 |
| **JMB2002 (Ab2001.08 N297A)** | FcγRI | 100 |
| | FcγRIIA R167 | Weak binding |
| | FcγRIIA H167 | |
| | FcγRIIIA F176 | |
| | FcγRIIIA V176 | |

**Table S3.** Binding of Ab2001.08 and JMB2002 to FcγRs was determined by BLI.

JM 036127
