## Supplementary material for "A human antibody with blocking activity to RBD proteins of multiple SARS-CoV-2 variants including B.1.351 showed potent prophylactic and therapeutic efficacy against SARS-CoV-2 in rhesus macaques": Table S4

JM 036127

**This PDF file includes:**

Fig. S1
Tables S1 to S4
References (39)

**Table S4A.**

| SARS-CoV-2 RBD proteins | Vendor | JMB2002 affinity $K_D$ (nM) | JMB2002 blocking activity $IC_{50}$, µg/ml （mean ± SD) | $IC_{50}$, nM （mean ± SD) |
|---|---|---|---|---|
| **SARS-CoV-2 RBD prototype** | ACRO | 3.33 | 0.102 ± 0.018 | 0.71 ± 0.13 |
| **SARS-CoV-2 RBD (N354D)** | | 1.44 | 0.087 ± 0.015 | 0.60 ± 0.10 |
| **SARS-CoV-2 RBD (V367F)** | | 2.10 | 0.058 ± 0.024 | 0.40 ± 0.17 |
| **SARS-CoV-2 RBD (W436R)** | | 2.32 | 0.024 ± 0.004 | 0.17 ± 0.03 |
| **SARS-CoV-2 RBD (R408I)** | | 1.42 | 0.085 ± 0.007 | 0.58 ± 0.05 |
| **SARS-CoV-2 RBD (N354D/D364Y)** | | 11.7 | 0.060 ± 0.012 | 0.41 ± 0.08 |
| **SARS-CoV-2 RBD (G476S)** | | 1.76 | 0.065 ± 0.002 | 0.45 ± 0.02 |
| **SARS-CoV-2 RBD (V483A)** | | 5.49 | 0.204 ± 0.057 | 1.41 ± 0.39 |
| **SARS-CoV-2 RBD (A475V)** | | 6.22 | 0.445 ± 0.112 | 3.07 ± 0.76 |
| **SARS-CoV-2 RBD (L452R)** | | 218 | 4.295 ± 2.023 | 28.63 ± 13.49 |

**Table S4A.** Binding affinity of JMB2002 to the SARS-CoV-2 RBD prototype and its variants was measured by BLI, and the blocking of SARS-CoV-2 RBD proteins to hACE2-Fc by JMB2002 was determined using ELISA.

JM 036127

**Table S4B.**

| SARS-CoV-2 S1 proteins | Vendor | JMB2002 affinity $K_D$ (nM) | JMB2002 blocking activity $IC_{50}$, µg/ml （mean ± SD) | $IC_{50}$, nM （mean ± SD) |
|---|---|---|---|---|
| **SARS-CoV-2 S1 prototype** | ACRO | 10.8 | 0.058 ± 0.019 | 0.40 ± 0.13 |
| **SARS-CoV-2 S1 (D614G)** | | 11.9 | 0.033 ± 0.007 | 0.23 ± 0.05 |
| **SARS-CoV-2 S1 UK mutant (H69del, V70del, Y144del, N501Y, A570D, D614G, P681H)** | Sino Biological | 5.70 | 0.995 ± 0.615 | 6.87 ± 4.24 |
| **SARS-CoV-2 S1 South African mutant (K417N, E484K, N501Y, D614G)** | | 1.59 | 0.165 ± 0.018 | 1.14 ± 0.12 |

**Table S4B.** Binding affinity of JMB2002 to the SARS-CoV-2 S1 prototype and its variants was measured by BLI, and the blocking of SARS-CoV-2 S1 proteins to hACE2-Fc by JMB2002 was determined using ELISA.

JM 036127
